# Demographic history impacts stratification in polygenic scores

**DOI:** 10.1101/2020.07.20.212530

**Authors:** Arslan A. Zaidi, Iain Mathieson

## Abstract

Large genome-wide association studies (GWAS) have identified many loci exhibiting small but statistically significant associations with complex traits and disease risk. However, control of population stratification continues to be a limiting factor, particularly when calculating polygenic scores where subtle biases can cumulatively lead to large errors. We simulated GWAS under realistic models of demographic history to study the effect of residual stratification in large GWAS. We show that when population structure is recent, it cannot be fully corrected using principal components based on common variants—the standard approach—because common variants are uninformative about recent demographic history. Consequently, polygenic scores calculated from such GWAS results are biased in that they recapitulate non-genetic environmental structure. Principal components calculated from rare variants or identity-by-descent segments largely correct for this structure if environmental effects are smooth. However, even these corrections are not effective for local or batch effects. While sibling-based association tests are immune to stratification, the hybrid approach of ascertaining variants in a standard GWAS and then re-estimating effect sizes in siblings reduces but does not eliminate bias. Finally, we show that rare variant burden tests are relatively robust to stratification. Our results demonstrate that the effect of population stratification on GWAS and polygenic scores depends not only on the frequencies of tested variants and the distribution of environmental effects but also on the demographic history of the population.

## Introduction

Population structure refers to patterns of genetic variation that arise due to non-random mating. If these patterns are correlated with environmental factors, they can lead to spurious associations and biased effect size estimates in genome-wide association studies (GWAS). Approaches such as genomic control (GC) [1], principal component analysis (PCA) [2], linear mixed models (LMMs) [3, 4] and linkage disequilibrium score regression (LDSC) [5] have been developed to detect and correct for this stratification. However, these approaches do not necessarily remove all stratification, particularly when multiple studies are meta-analyzed [6, 7]. Large GWAS in relatively homogeneous populations, such as the UK Biobank (UKB) [8], should alleviate many of these concerns. However, such populations still exhibit fine-scale population structure [9–15]. The extent to which this fine structure impacts GWAS inference in practice is largely unknown, and it is not clear whether existing methods adequately correct for it. This question has become increasingly acute in light of the recent focus on polygenic scores for disease risk prediction [16, 17]. Polygenic scores for many physical and behavioral traits exhibit geographic clustering within the UK even after stringent correction for population structure with standard methods [12, 18]. While some of this variation may be attributed to recent migration patterns [18], it could also reflect residual stratification in variant effect sizes [19].

To address these questions, we investigate the effect of population structure on GWAS in a simulated population with a similar degree of structure to the UK Biobank. We consider the fact that different demographic histories can give rise to the same overall degree of population structure (in terms of statistics such as *F*_*ST*_ and the genomic inflation factor, *λ*). This is relevant because the degree to which common and rare variants are impacted by, and are thus informative about, population structure depends on the demographic history of the population. It is therefore important to understand the demographic history of GWAS populations in order to understand the consequences of stratification. To demonstrate this, we leveraged recent advances in our understanding of human history to simulate GWAS under different realistic demographic models.

## Results

### Rare variants capture recent population structure

We simulated population structure using a six-by-six lattice-grid arrangement of demes with two different symmetric stepping-stone migration models (Fig. 1). First, a model where the structure extends infinitely far back in time (perpetual model; e.g. [20]), and second, a model where the structure originated 100 generations ago (recent structure model). This second model is motivated by the observation from ancient DNA that Britain experienced an almost complete population replacement within the last 4,500 years [21], providing an upper bound for the establishment of present-day geographic structure in Britain. We set the migration rates in the two models to match the degree of population structure in the UK Biobank, measured by the average *F*_*ST*_ between regions [9] and the genomic inflation factor for a GWAS of birthplace in individuals with ‘White British’ ancestry from the UK Biobank [12] (Methods).

**Figure 1:**
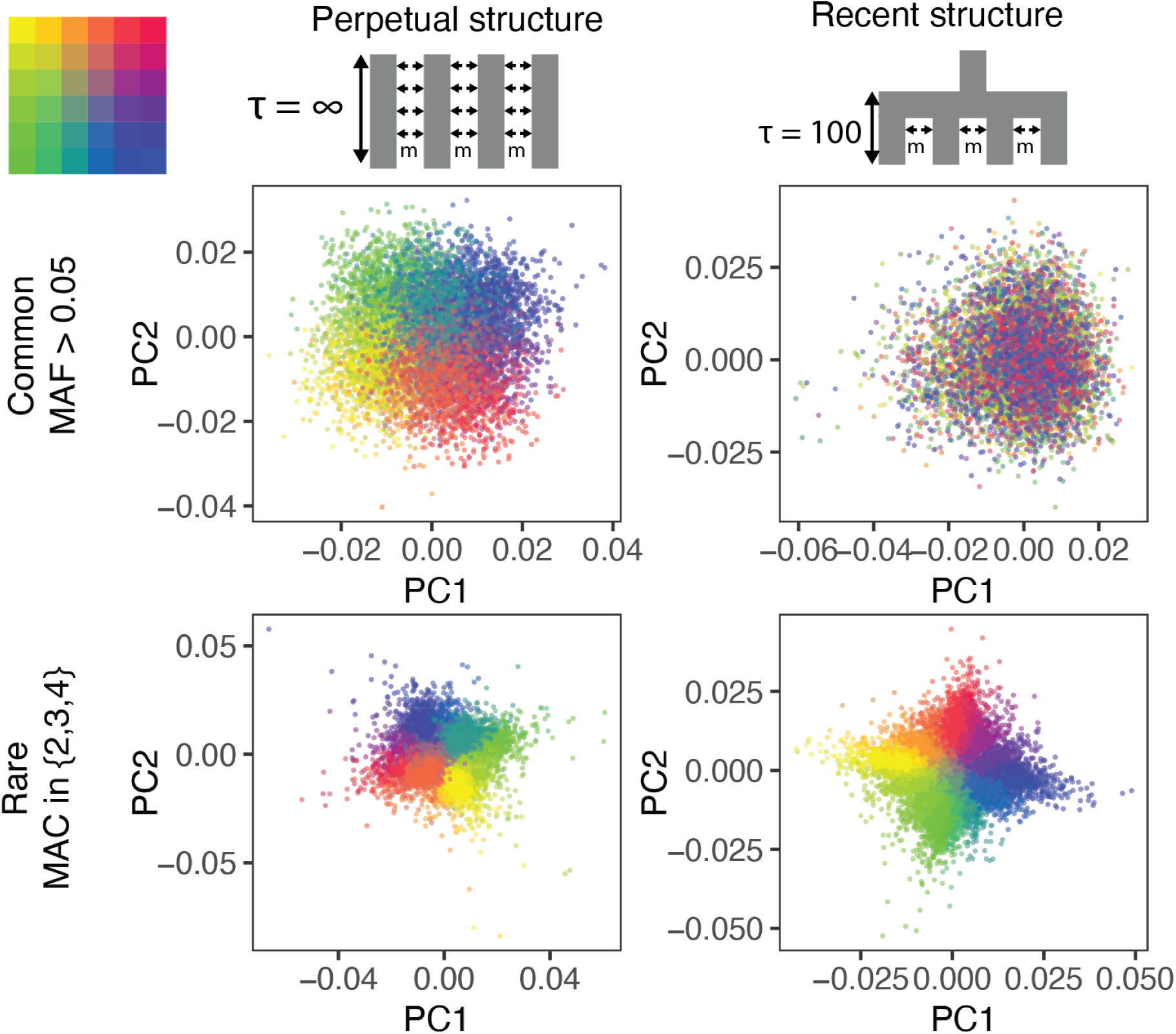
The ability of PCA to capture population structure depends on the frequency of the variants used and the demographic history of the sample. Panels show the first and second principal components of the genetic relationship matrix constructed from either common (upper row) or rare (lower row) variants. Each point is an individual (N = 9,000) and their color represents the deme in the grid (upper left) from which they were sampled. Both common (minor allele frequency >0.05) and rare (minor allele count=2,3 or 4) variants can be informative when population structure is ancient (left column; *τ* = *∞* represents the time in generations in the past at which structure disappears) but only rare variants are informative about recent population structure (right column; *τ* = 100 generations). Number of variants used for PCA: 200,000 (upper row), 1 million (lower left), and *≈*750,000 (lower right).

The two models produce qualitatively different population structure, even though *F*_*ST*_ is the same. When structure is recent, it is driven largely by rare variants which tend to have a more recent origin [22–24] and are therefore less likely to be shared among demes. Common variants, because they are older and usually predate the onset of structure in our model, are more likely to be shared among demes and have not drifted enough in 100 generations to capture the spatial structure effectively. Therefore recent structure is captured by the principal components of rare variants (rare-PCA) but not common variants (common-PCA) (Fig. 1). In comparison, when population structure is perpetual, both common and rare variants carry information about spatial structure (Fig. 1). When rare variants are unavailable (for example, from genotype array data), PCA of haplotype or identity-by-descent (IBD) sharing is similarly informative about recent population structure (Fig. S1).

### The impact of population stratification depends on demographic history

That common variants fail to capture recent population structure has important implications for GWAS. Most GWAS use PCA or LMMs, both of which rely on the genetic relatedness matrix (GRM) to describe population structure. Since rare variants are not well represented on SNP arrays, the GRM is usually constructed from common variants. This will lead to insufficient correction if common variants do not adequately capture recent population structure. To test this, we simulated a GWAS (N=9,000) of a non-heritable phenotype (i.e. *h*^2^ = 0) with an environmental component that is either smoothly (e.g. latitude) or sharply (e.g. local effects) distributed in space (Fig. 2; Methods). We calculated GRMs using either common (minor allele frequency, MAF > 0.05) or rare variants (minor allele count, MAC=2, 3, or 4), and included the first 100 PCs in the model to correct for population structure.

**Figure 2:**
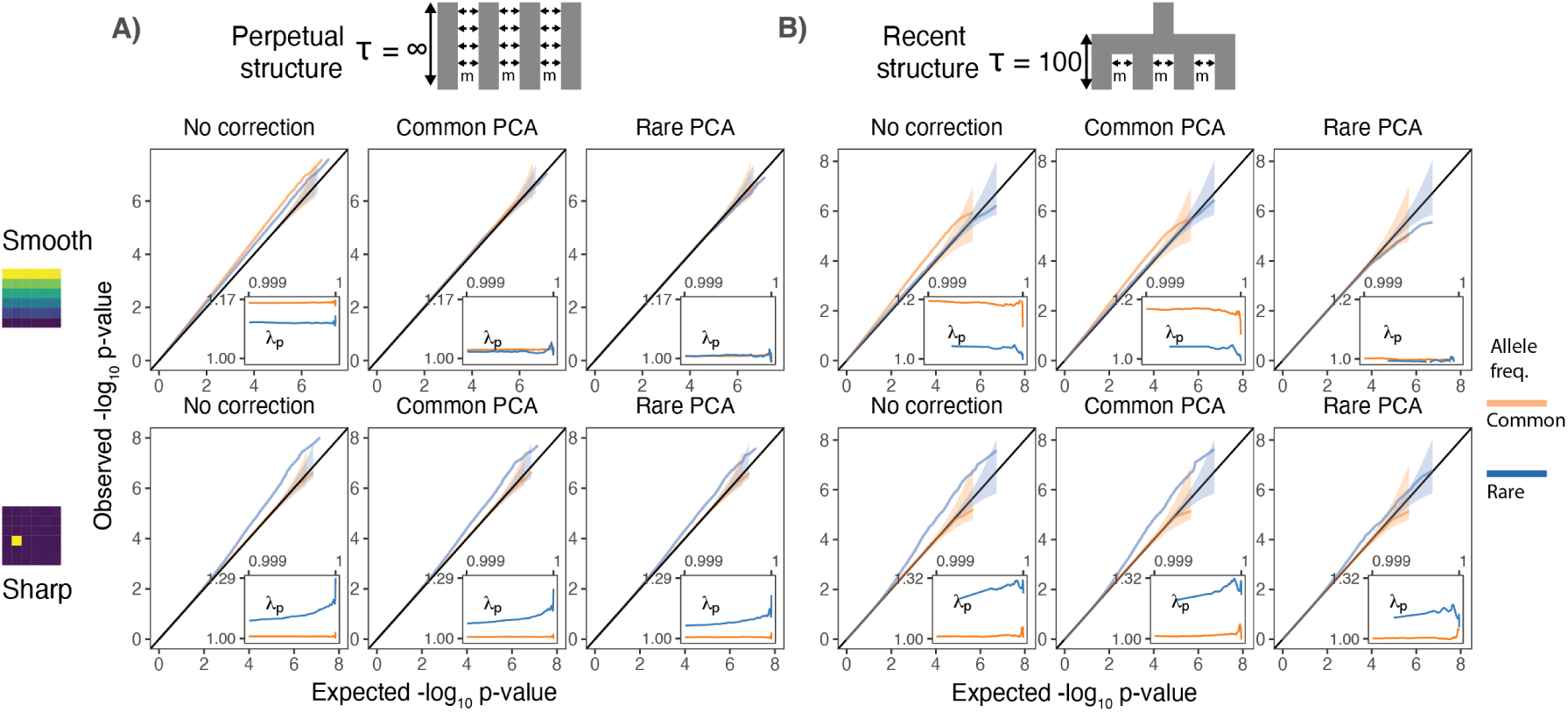
Test statistic inflation under two different demographic histories: A) perpetual and B) recent structure. Upper and lower rows show results for smoothly and sharply distributed environmental risk respectively, while columns show different methods of correction. The simulated phenotype has no genetic contribution so any deviation from the diagonal represents inflation in the test statistic. Each panel shows QQ plots for -log_10_ p-value for common (orange) and rare (blue) variants. Insets show inflation (*λ*_*p*_) in the tail (99.9^th^ percentile) of the distribution. Results are averaged across 20 simulations of the phenotype.

When population structure is recent, smooth environmental effects lead to an inflation in common, but not rare, variants and this inflation can only be corrected with rare-but not common-PCA (Fig. 2B, top row). This is a consequence of the fact that rare variants carry more information about recent structure than common variants (Fig. 1). In contrast, under the perpetual structure model, both common and rare variants may be inflated due to smooth environmental effects (Fig. 2A, top row), but this inflation is largely corrected with either common- or rare-PCA (Fig. 2A, top row).

In the perpetual structure model, local environmental effects only impact rare variants [20] (Fig. 2A, lower row). But, when the structure is recent, local effects may confound even common variants (Fig. 2B, lower row). The inflation in rare variants due to local effects cannot be fully corrected using either common- or rare-PCA (Fig. 2A and B, lower row) because the environmental effects cannot be represented by a linear combination of the first hundred principal components. Importantly, local effects only impact a small subset of variants—those clustered in the affected deme(s)—resulting in inflation only in the tails of the test statistic distribution (Fig. 2). This pattern of inflation cannot be detected using standard genomic inflation, which assumes that stratification impacts enough variants to shift the median of the test statistic [1], making it difficult to distinguish between true associations and residual stratification. We find similar results when we use LMMs instead of PCA (Fig. S2). This is expected, since both rely on the same construction of the GRM. Therefore, in large studies with recent structure, such as the UKB, neither PCA-nor LMM-based methods will fully correct for stratification as long as the GRM is derived from common variants.

### Polygenic scores capture residual environmental stratification

Polygenic scores—constructed by summing the effects of large numbers of associated variants— offer a simple way to make genetic risk predictions. At least in European ancestry populations, they can explain a substantial proportion of the phenotypic variance in complex traits like height [25], BMI [25], and coronary artery disease risk [26]. However, their practical utility is limited by lack of transferability between populations [27] and between subgroups within populations [28]. This may be due in part to residual stratification in polygenic scores. To understand the behavior of polygenic scores under the perpetual and recent structure models, we simulated GWAS (N=9,000) of a heritable phenotype with a genetic architecture similar to that of height. We used GWAS effect sizes to calculate polygenic scores in an independent sample (N=9,000) and subtracted the true genetic values for each individual to examine the spatial bias in polygenic scores due to stratification.

Under both perpetual and recent structure models, residual polygenic scores are spatially structured, recapitulating environmental effects even when 100 common-PCs are used as covariates in the GWAS (Fig. 3). LMMs perform similarly (Fig. S3). This is due to the fact that when population stratification is not fully corrected, the effect sizes of variants that are correlated with the environment tend to be over or underestimated depending on the direction and strength of correlation (Fig. S4). Stratification in residual polygenic scores is minimal when the causal variants are known, but not when the score is constructed from the most significant SNPs (‘lead SNPs’) (Fig. 3)—almost always the case in practice. Picking the most significant SNPs tends to enrich for variants that are more structured than the causal variants. Thus, polygenic scores are especially prone to residual stratification when constructed using SNPs in ‘sub-significant’ peaks where the effect of stratification dominates over relatively small causal variant effect sizes.

**Figure 3:**
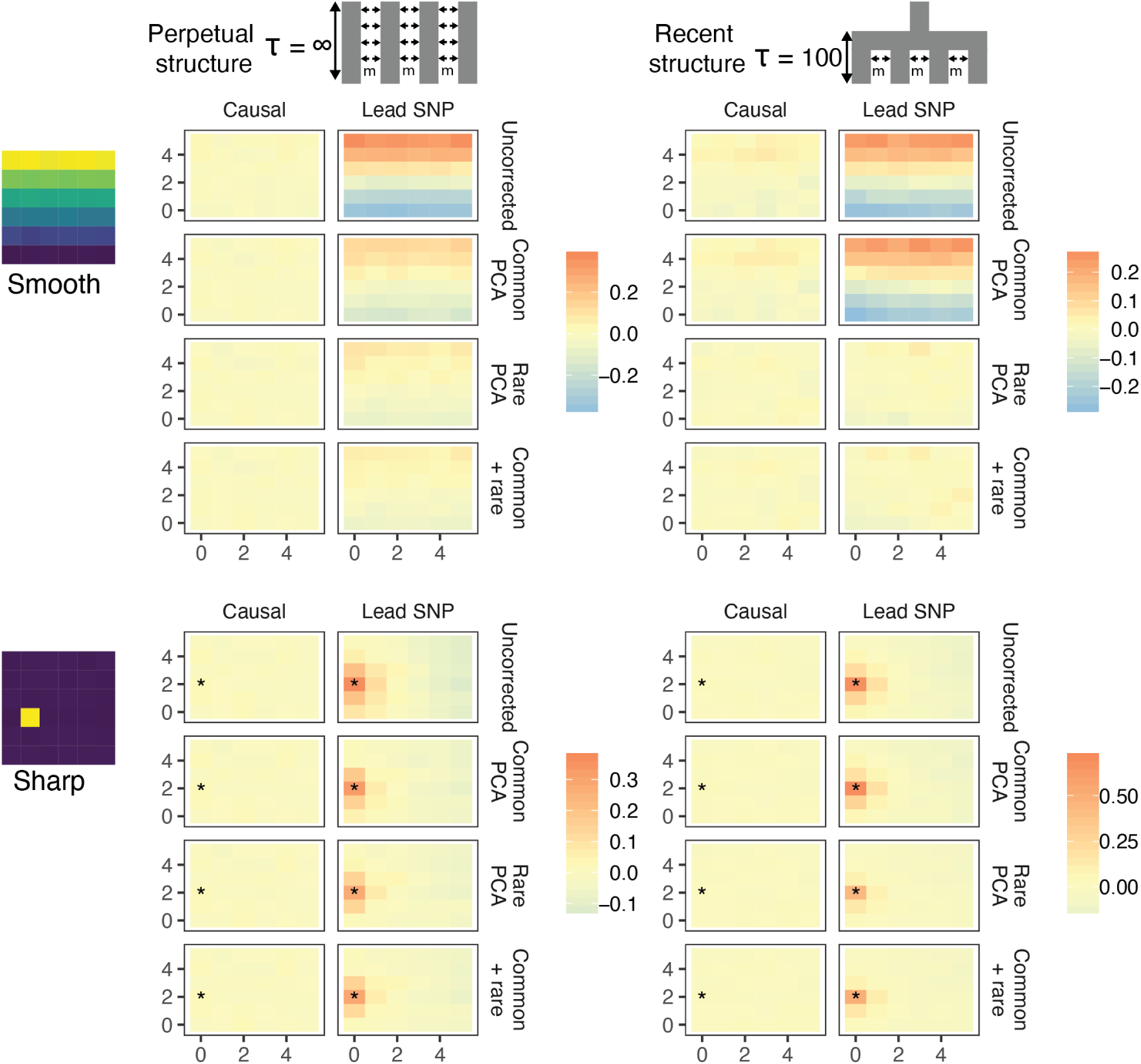
Residual stratification in effect size estimates translates to residual spatial structure in the polygenic score in the (A) recent and (B) perpetual structure models. The simulated phenotype in the training sample has a heritability of 0.8, distributed over 2,000 causal variants. Each small square is colored with the mean residual polygenic score for that deme in the test sample, averaged over 20 independent simulations of the phenotype. In each panel, the rows represent different methods of PCA correction and columns represent two different methods of variant ascertainment. ‘Causal’ refers to causal variants with p-value < 5×10^−4^, and ‘lead SNP’ refers to a set of variants, where each represents the most significantly associated SNP with a p-value < 5×10^−4^ in a 100kb window around the causal variant. The simulated environment is shown on the left. For the sharp effect, the affected deme is highlighted with an asterisk.

### The effect of stratification in more complex models

In reality, genetic structure in most studies is more complex than either model discussed above. Most populations are genetically heterogeneous and each genome is shaped by processes such as ancient and recent admixture, non-random mating and selection, all of which vary both spatially and temporally. The present-day population of Britain, for example, is the result of a complex history of admixture and migration [9, 21]. Thus, restricting analysis even to the ‘White British’ subset of UK Biobank involves population structure on multiple time scales. To study these effects, we simulated under a model based on the demographic history of Europe and geographic structure of England and Wales, while maintaining the same degree of structure as the previous models (Fig. 4). In addition to recent geographic structure, we simulated an admixture event 100 generations ago between two populations, each of which are themselves the result of mixtures between many ancient populations (Fig. 4). We varied the admixture fraction from the two source populations to create a North-South ancestry cline and sampled individuals to mimic uneven sampling in the UK Biobank (Fig. 4, Methods).

**Figure 4:**
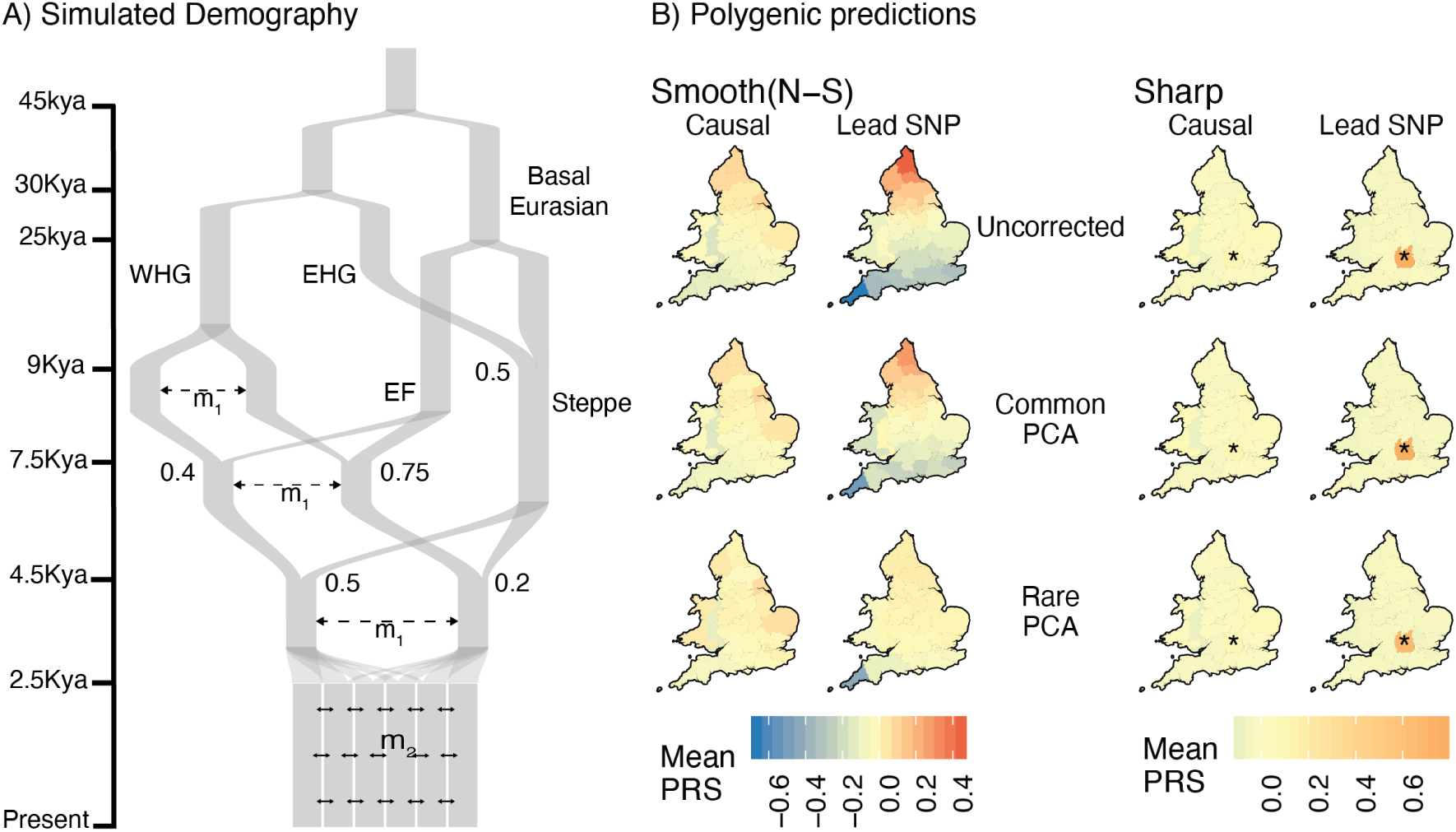
Residual stratification in polygenic scores under a complex demographic model, geographic structure representing England and Wales, and non-uniform sampling. (A) Il-lustration of the simulated demography. (B) Maps depicting the spatial distribution of residual polygenic scores, as in Fig. 3, averaged across 20 simulations of the phenotype. Columns: ‘Smooth’ and ‘sharp’ refers to the environmental effect and ‘causal’ and ‘lead SNP’ refer to sets of variants that were used to construct polygenic scores. Rows: Different methods of correction for population structure.(WHG and EHG: Western and Eastern Hunter Gatherers; EF: Early Farmers)

The results under this model are very similar to the recent structure model in that when the environmental effect is smoothly distributed, it cannot be corrected using common-PCA as population structure is largely recent (Fig. 4). Note also that correction is not complete even with rare-PCA as seen from the biased polygenic scores of individuals from Cornwall. This is not due to reduced migration in the region (‘edge effects’) but rather to uneven sampling (only 17 individuals sampled from Cornwall as opposed to 250 under uniform sampling). The bias disappears when individuals are sampled uniformly (Fig. S6). Thus, our ability to correct for stratification and the utility of polygenic scores also depends on the sampling design of the GWAS. As with the other models, local effects cannot be corrected using either common- or rare-PCA (Fig. 4).

### Sibling-based tests are robust to environmental stratification

Sibling-based studies test for association between the phenotypic difference between siblings and the difference in their genotypes. These, and other family-based association tests, are robust to population stratification as any difference in siblings’ genotypes is due to Mendelian segregation and therefore uncorrelated with environmental effects. We simulated sibling pairs under the recent structure model and confirmed that polygenic scores constructed using SNPs and their effect sizes from the sibling-based tests were uncorrelated with environmental variation (Fig. 5 lower row).

**Figure 5:**
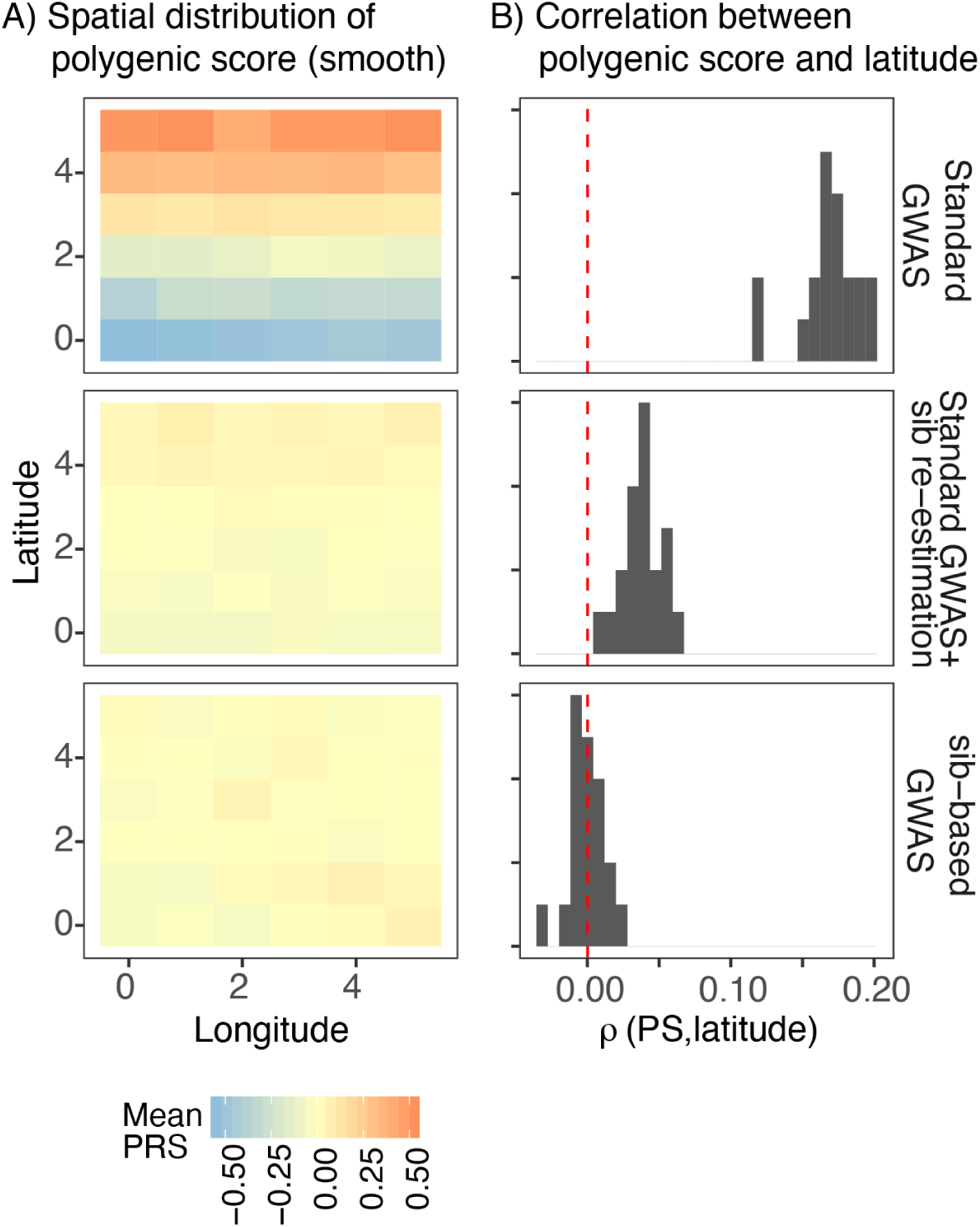
Comparison of stratification in polygenic scores between standard and sibling-based association tests under the recent structure model. Phenotypes simulated as in Fig. 3. (A) Spatial distribution of polygenic scores generated using (top) effects of variants detected through standard GWAS, (middle) variants ascertained in standard GWAS but with effect sizes re-estimated in sib-based design, (bottom) variants ascertained and effect sizes estimated in sib-based design. In each case, the effect is averaged over 20 simulations. (B) Distribution of correlation between polygenic score and latitude for 20 simulations of the smooth environmental effect.

In practice, however, sample sizes for sibling-based studies are much smaller than standard GWAS. A possible hybrid approach is to first ascertain significantly associated SNPs in a standard GWAS and then re-estimate effect sizes in siblings. However, this is not completely immune to stratification. To demonstrate, we took the significant lead SNPs from a standard GWAS, re-estimated their effect sizes in an independent set of 9,000 sibling pairs simulated under the same demographic model, and then generated polygenic scores in a third, independent, sample of 9,000 unrelated individuals. Polygenic scores generated this way are still correlated with the environmental effect when it is smoothly distributed, although less than when effects sizes from the standard GWAS are used (Fig. 5). Even though the sibling re-estimated effects are unbiased, stratification in the polygenic score persists because the frequencies of the lead SNPs are preferentially correlated with the environment. This is less pronounced for local effects because stratification is driven by variants that are rare in the discovery sample and often absent in the test sample (Fig. S5).

### Burden tests are relatively robust to local environmental effects

Finally, we studied the effect of population stratification on burden tests, which aggregate rare variants in a gene or other functional category to circumvent the limited statistical power of single rare variant tests. Specifically, we examined the behavior of a simple burden statistic—the total number of rare derived alleles (frequency < 0.001) in each gene. While burden tests have the potential to be affected by rare variant stratification [20], we find that with our parameters, for an average sized gene (total exon length of *≈* 1.3kb, mean of 16 rare variants), burden tests are robust to environmental effects under both perpetual and recent structure models (Fig. 6). Because the rare variant burden involves averaging over many variants, the burden statistic behaves more like a common variant than a rare variant in terms of its spatial distribution (Fig. S7). Consequently, it is more susceptible to confounding by smoothly distributed environmental effects and relatively immune to local effects (Fig. 6). Similar to common variants, rare-PCA corrects the inflation when the environment is smooth (Fig. 6). More generally, the spatial distribution of gene burden depends on the number of variants and genetic distance across which it is aggregated. Gene burden should become geographically less localized with an increase in the number of aggregated rare variants as each is likely to arise in an independent branch of the genealogy (Fig. S7). As genetic distance between mutations increases, recombination decouples genealogies, reducing the probability of multiple mutations arising on the same branch. Thus, the rare variant burden in short genes with little recombination behaves more like a rare variant and is susceptible to local effects (Fig. 6B lower row) just as single rare variants.

**Figure 6:**
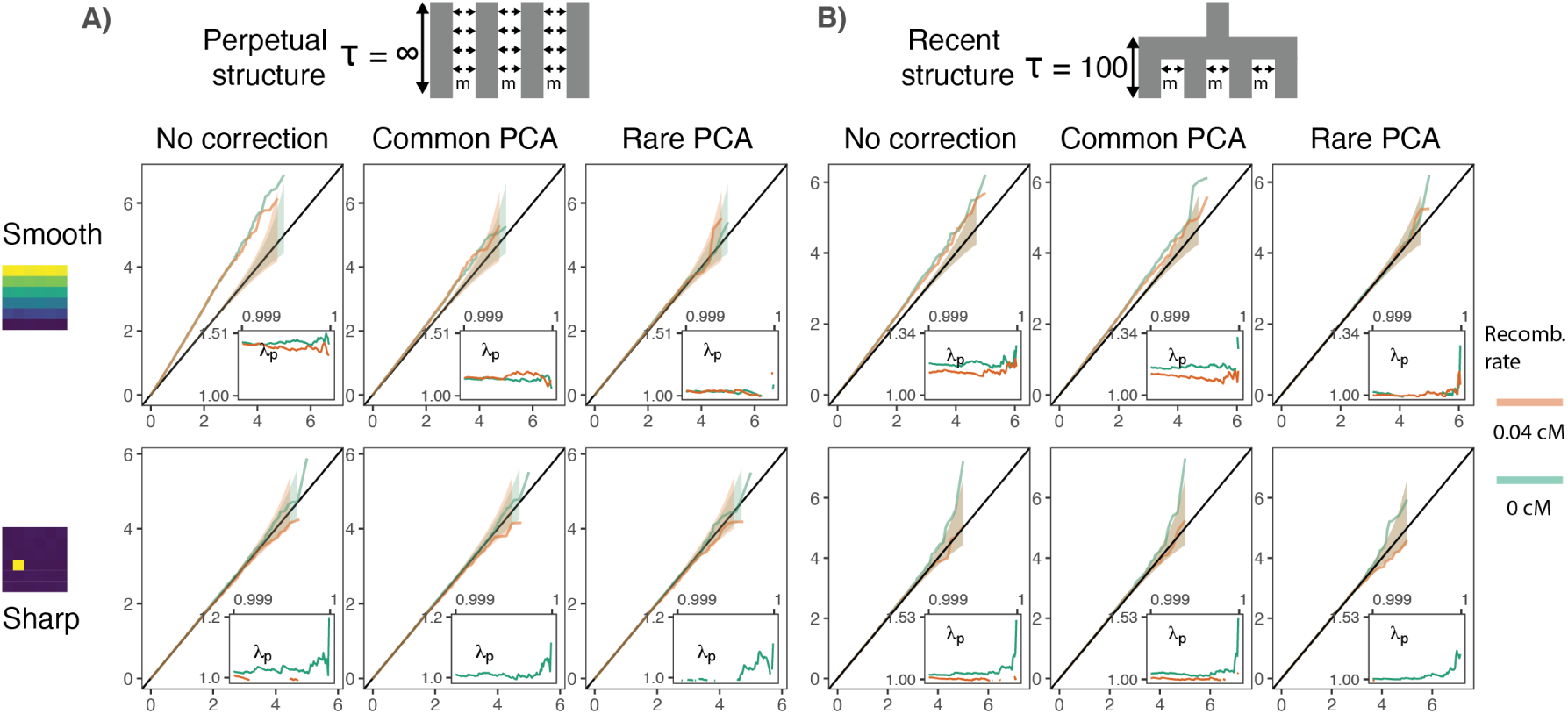
Gene burden tests are relatively robust to stratification. QQ plots of expected and observed -log_10_ P-value under the (A) perpetual and (B) recent structure models for the association of rare variant burden across a gene with total exon length of 1.3 kb and non-heritable phenotype with a smooth (upper) or sharp (lower) distribution of environmental effects. Orange and green lines show results for a gene with and without recombination, respectively. Inset shows inflation in the tail (99.9%) of the test statistic distribution.

## Discussion

The effect of population structure on GWAS depends on the amount of structure, the frequency of the variants tested and the distribution of confounding environmental effects. Here, we demonstrated that it also depends on the demographic history of the population in a way that is not fully captured by the degree of structure as summarized by *F*_*ST*_ and genomic inflation. Consequently, to fully correct for population structure, it is necessary to know not only the degree of realized structure, but also the demographic history that generated it, and the likely distribution of environmental effects.

Generally, PCA (or mixed models) based on common variants will inadequately capture and correct population structure with a recent origin. This might partly explain why polygenic scores derived from the UK Biobank exhibit geographic clustering [12, 18]. In such cases, PCA based on rare variants would be more effective at controlling for fine structure, as rare variants are more informative about recent population history [22–24, 29, 30]. Haplotype sharing [31] or identity-by-descent (IBD) segments are similarly informative about recent history [32, 33], and provide an alternative to rare variant PCA when sequence data are not available, or when there are relatively few rare variants to adequately capture the structure, for example in exome-sequence data. However, this still leaves the question of exactly what frequency of rare variants (or length of IBD segments) to use. Since structure in most populations exists on multiple time scales, the best approach might be to use multiple sets of PCs derived from variants across the frequency spectrum.

This approach would correct for environmental effects that are relatively smoothly distributed with respect to ancestry and can be expressed as a simple linear function of the principal components. Sharply distributed effects (e.g. local environment or batch effects) may not be fully corrected with any method, regardless of the demographic history of the population. Local effects are an important concern particularly for rare variants and large studies where even subtle effects will significantly impact effect size estimates. Because local effects lead to inflation in the tail of the test statistic, rare variant associations should always be treated with caution. Rare variant burden tests are more robust to local effects than single variant tests, but burden statistics may be heterogeneous in their response to stratification. For example, short genes are more affected by recent structure than long genes.

Even imperfect correction for population structure is probably suficient to limit the number of genome-wide false positive associations when GWAS are used for mapping. But when large numbers of marginally associated variants are used to construct polygenic scores, even small overestimates in effect sizes can lead to substantial bias. Essentially some of the predictive power of the polygenic score will derive from predicting environmental structure rather than genetic effects. Comparison of polygenic scores derived from standard GWAS and sibling-based studies suggests that this effect can be substantial [28], and it may also contribute to inflated estimates of heritability and genetic correlation. Even though family-based studies are immune to stratification, we show that the practice of discovering associations in a standard GWAS and then re-estimating their effects in siblings can introduce stratification in polygenic scores if there is inadequate correction in the original GWAS.

Our study focused on population structure arising from ancient admixtures and geographic structure because these are relatively well-understood and easy to model. However, our results generalize to any type of population structure, for example due to social stratification or assortative mating. What we refer to as local environmental effects also includes socially structured factors like cultural practices. Ultimately, no single approach can completely correct for population stratification. Empirical approaches including PCA based on variants across the frequency spectrum or IBD sharing, and inclusion of geographic and other covariates in GWAS analyses will help. Replication in within-family studies and populations of different ancestry provide greater confidence while simulations informed by demographic history can help to check robustness and provide a quantitative estimate of bias.

## Methods

### Simulations of population structure

We used *msprime* [34] to simulate genotypes in a 6×6 grid of demes and modeled the demoraphic history in three different ways: (i) where the structure extends infinitely far back in time (‘perpetual’), (ii) where all demes collapse into a single population 100 generations in the past (‘recent’), and (iii) a more complex model that is loosely based on the demographic history of Europe ([35]) (Fig. 4; ‘complex’). The effective population size of all demes and we fixed the merged ancestral population size to 10,000 diploid individuals.

For the perpetual and recent models, we parameterized the degree of structure in the data with a fixed, symmetric migration rate among demes (*m*) chosen to match the degree of structure observed in Britain. To select an appropriate value for *m*, we simulated a 10Mb genome (10 chromosomes of 1Mb each) with mutation and recombination rates of 1× 10^−8^ per-base per-generation, for 9,000 individuals (250 per-deme) for a range of migration rates under each demographic model (Table S1). We estimated mean *F*_*ST*_ across all demes with the Weir and Cockerham estimator [36] using an LD-pruned (PLINK –indep-pairwise 100 10 0.1 [37, 38]) set of common variants (MAF > 0.05). We used the ratio of averages approach [39] to calculate *F*_*ST*_ and estimated genomic inflation on birthplace (*λ*_*location*_) by carrying out GWAS on an individual’s x and y coordinates in the grid, similar to the GWAS on longitude and latitude in [12]. The migration rate was chosen for each model separately to roughly match the mean *F*_*ST*_ observed among regions in Britain (*≈* 0.0007) [9] and *λ*_*location*_ *≈* 12 reported for the UKB [12]. Because genomic inflation scales linearly with sample size [40], we matched the expected value given our sample size of 9K using:

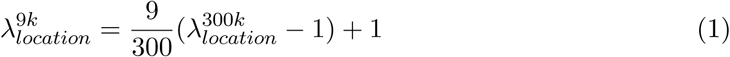

Where 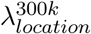 is the observed value (*≈* 12) given a sample size of 300,000 as in Ref. [12]. Plugging this in, we get an expected value of 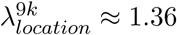. To match this approximately, we set the migration rate to a fixed value of 0.05 and 0.07 for the ‘recent’ and ‘perpetual’ models, respectively (Table S1).

We parameterized the ‘complex’ model with two migration rates, *m*_1_ and *m*_2_, where *m*_1_ represents the migration rate between the source populations mixing 100 generations before present (2.5Kya) and *m*_2_ represents the migration rate between adjacent demes in the grid (Fig. 4). We selected *m*_1_ and *m*_2_ in a step-wise manner, first setting *m*_1_ = 0.004 (representing the *F*_*ST*_ between the two source populations) to match the maximum *F*_*ST*_ between regions in Britain. We then set *m*_2_ = 0.08 (representing subsequent mixing and isolation by distance) to match the mean *F*_*ST*_ between regions in Britain [9] (Table S1). In all cases, after selecting the appropriate migration parameters, we re-simulated genotypes under each model for a larger genome of 200Mb (20 chromosomes of 10Mb each) which we used for all further analysis.

### Geographic structure in England and Wales

We downloaded the Nomenclature of Territorial Units for Statistics level 2 (NUTS2) map for 35 regions in England and Wales (version 2015) from data.gov.uk and assigned each individual of ‘White British’ ancestry in the UKB to a region based on their birthplace. We calculated the proportion of individuals sampled from each region and used these as weights in our simulations to mimic the sampling distribution in the UKB. To generate a migration matrix between regions, we generated an adjacency matrix for the NUTS2 districts using the ‘simple features’ (sf) R package [41], where an entry is one if two districts abut and zero otherwise, and multiplied this matrix by the migration parameter *m*_2_.

### Simulation of phenotypes

To study the effect of stratification on test statistic inflation, we simulated non-heritable phenotypes *y*_*ij*_ of an individual *i* from deme *j* as *y*_*ij*_ *∼ N* (*µ*_*j*_, *σ*), where *µ*_*j*_ is the mean environmental effect in deme *j*. For the smooth effect, we chose *µ*_*j*_ such that the difference between the northern and southernmost demes was 2*σ*. For the sharp effect, we set *µ*_*j*_ = 2*s* for one affected deme and zero otherwise. To test the impact of population structure on effect size estimation and polygenic score prediction, we simulated heritable phenotypes using the model described in [42]. We selected 2,000 variants across the 200Mb genome (one variant chosen uniformly at random in each 100kb window) and sampled their effect sizes as 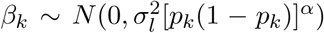 where 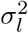 is the frequency-independent component of genetic variance, *p*_*k*_ is the allele frequency of the *k*^*th*^ variant, and *α* is a scaling factor. We set *α* = *-*0.4 based on an estimate for height [42] and 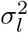 such that the overall genetic variance underlying the trait,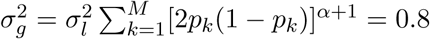. We calculated the genetic value for each individual, 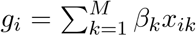, where *x*_*ik*_ is the number of derived alleles individual *i* carries at variant *k*, and added environmental effects as described above. We generated 20 random iterations of both heritable and non-heritable phenotypes.

### GWAS

We simulated 18,000 individuals (500 from each deme) under each demographic model and split the sample into two equally sized sets, a training set on which GWAS and PCA were carried out, and a test set for polygenic score predictions. To carry out common- and rare-PCA, we first sampled 200,000 common (MAF>5%) and one million rare (minor allele count = 2, 3, or 4) variants, respectively, from all variants generated under each model, and generated PCs using PLINK [38]. To carry out PCA on identity-by-descent (IBD) sharing, we called long (>10cM) pairwise IBD segments using GERMLINE [43] with default parameters and generated an IBD-sharing GRM, in which each entry represents the total fraction of the haploid genome (100Mb) shared by individual pairs. We calculated eigenvectors (PCs) of the IBD-sharing GRM using GCTA [44]. and performed GWAS using --glm in PLINK 2.0 with 100 PCs as covariates [38].

We fitted linear mixed models using GCTA-LOCO [44] where the GRM was based on the same common or rare variants used for PCA. GCTA’s LOCO (leave one chromosome out) algorithm fits a model where the GRM is constructed from SNPs that are not present on the same chromosome as the variant being tested to avoid proximal contamination. We also included the top 100 PCs as fixed effects in the mixed models.

We calculated genomic inflation (*λ*_*p*_) for non-heritable phenotypes as 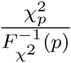 where 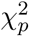 is the *p*^*th*^ percentile of the observed association test statistic and 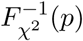 is the quantile function of the *χ*^2^ distribution with 1 degree of freedom.

### Sibling-based tests

We conducted structured matings by sampling pairs of individuals from the same deme and generated the haplotypes of each child by sampling haplotypes, with replacement, from each parent without recombination. We generated heritable phenotypes as described in the previous section for each sibling and modeled the effect of each variant as

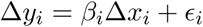

where Δ*y* is the difference in siblings’ phenotypic values, Δ*x*_*i*_ is the difference in the number of derived alleles at the *i*^*th*^ variant.

### Polygenic scores

We calculated polygenic scores for each individual as 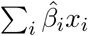 where 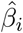 is the estimated effect size and *x*_*i*_ is the number of derived alleles for the *i*^*th*^ variant. Because we were interested in studying patterns of *residual* stratification, we subtracted individuals’ true (simulated) genetic values, which themselves can be structured, from polygenic scores. We averaged residual polygenic scores across 20 random iterations of causal variant selection, effect size generation, and GWAS to minimize stochastic variation.

### Gene burden

We simulated genes, each with eight exons of length 160 bp separated by introns of length 6,938 bp, representing an average gene in the human genome [45]. We simulated 100,0000 genes for the ‘recent’ model with and without recombination and for the ‘perpetual’ model with no recombination. For the ‘perpetual’ model with recombination, we simulated 50,000 genes. We calculated gene burden as the total count of derived alleles (frequency < 0.001) across all exons in the gene for each individual. For genes simulated without recombination, we only simulated the exons because the introns serve no purpose and add to the computational cost. To ensure that differences in structure in gene burden between models was driven by differences in demographic history and not differences in the number of rare variants, we first calculated the mean (16) and standard deviation (4) of the number of rare variants under the ‘recent’ model and sampled from this distribution when simulating under the ‘perpetual’ model. The geographic clustering of burden was measured using Gini curves and the Gini coeffcient.

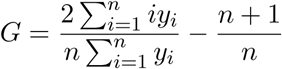

where *y*_*i*_ is the cumulative gene burden in the *i*^*th*^ deme sorted in increasing order of gene-burden and *n* is the number of demes. The Gini coefficient ranges from zero, indicating that the burden is uniformly distributed in space, to one, indicating that the burden is concentrated in a single deme (Fig. S7).

## Code availability

We carried out all analyses with code written in Python 3.5, R 3.5.1, and shell scripts, which are all available at https://github.com/Arslan-Zaidi/popstructure.

## Acknowledgments

This research was supported by NIGMS award number [R35GM133708]. The content is solely the responsibility of the authors and does not necessarily represent the offcial views of the National Institutes of Health. The UK Biobank Resource was used under Application 33923.

## Supplementary material

**Figure S1:**
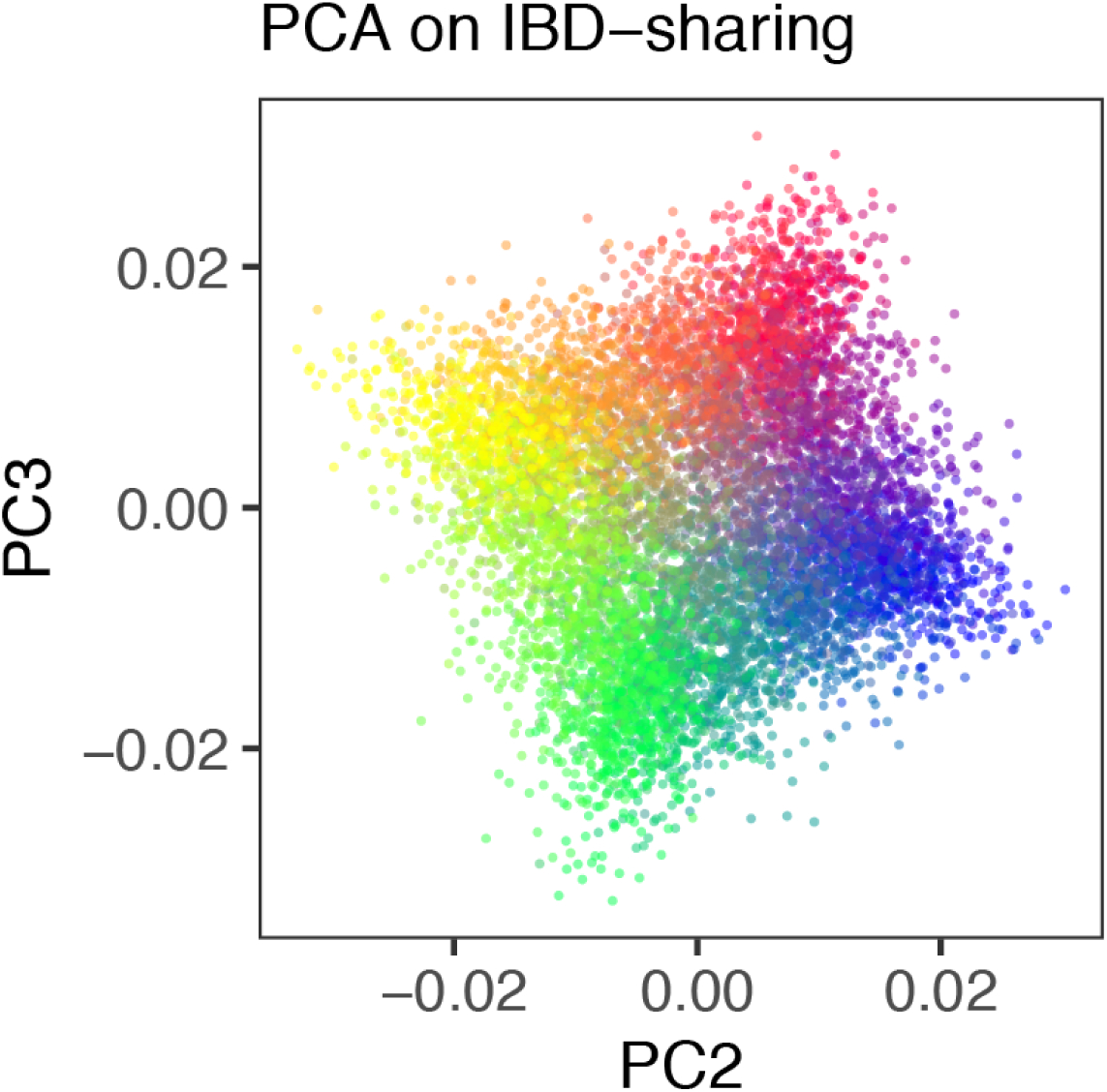
PCA based on Identity-by-descent sharing recapitulates fine structure under the recent structure model.

**Figure S2:**
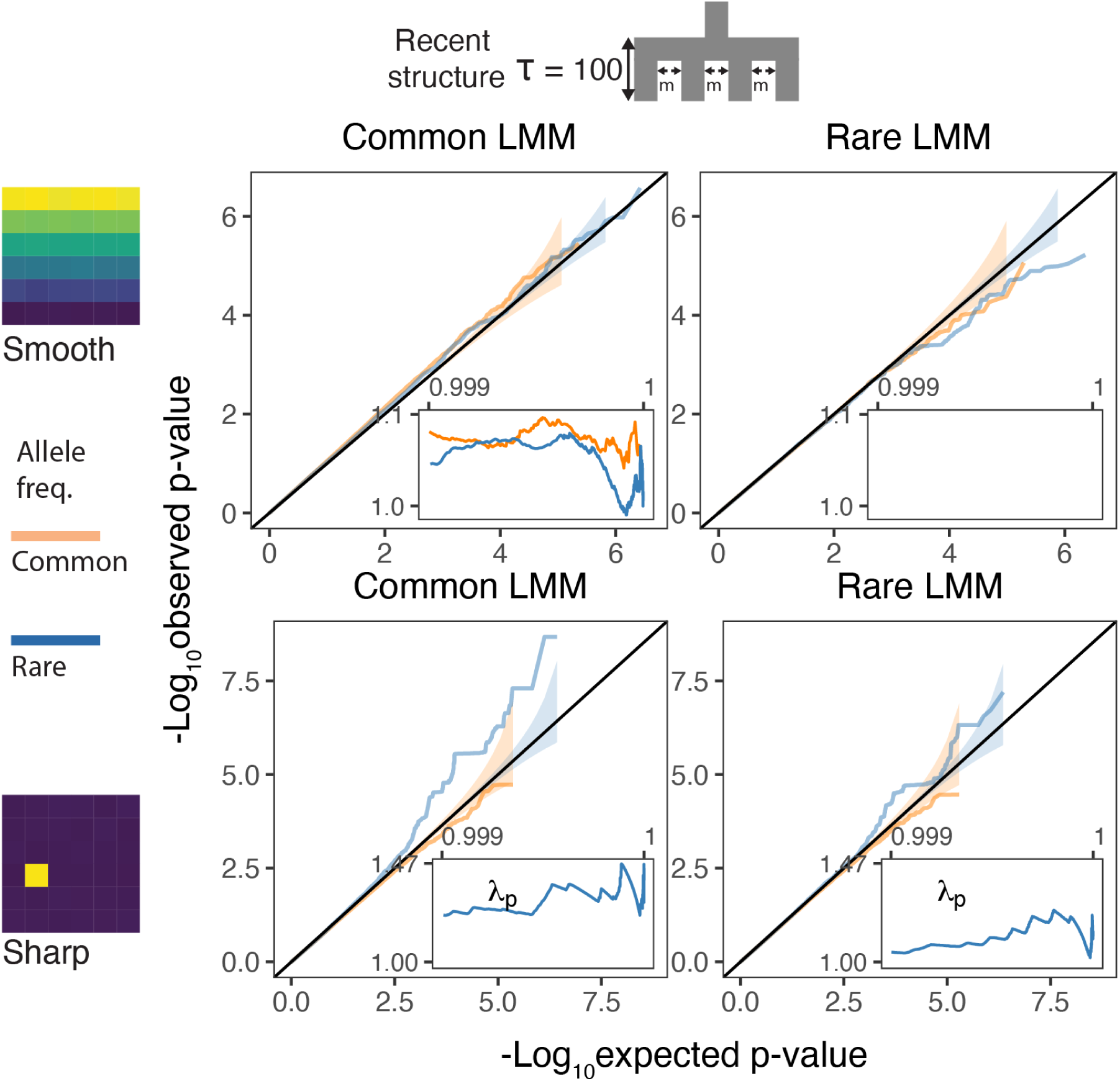
QQplots for linear mixed model association of non-heritable phenotypes carried out with GCTA-LOCO for genotypes generated under the recent structure model. Models were fit with either a GRM constructed with common variants (left column) or with rare variants (right column). Insets show the 99.9% tail of the inflation factor. Orange and blue colored lines refer to common and rare variants, respectively. Results shown for a single simulation of the phenotype.

**Figure S3:**
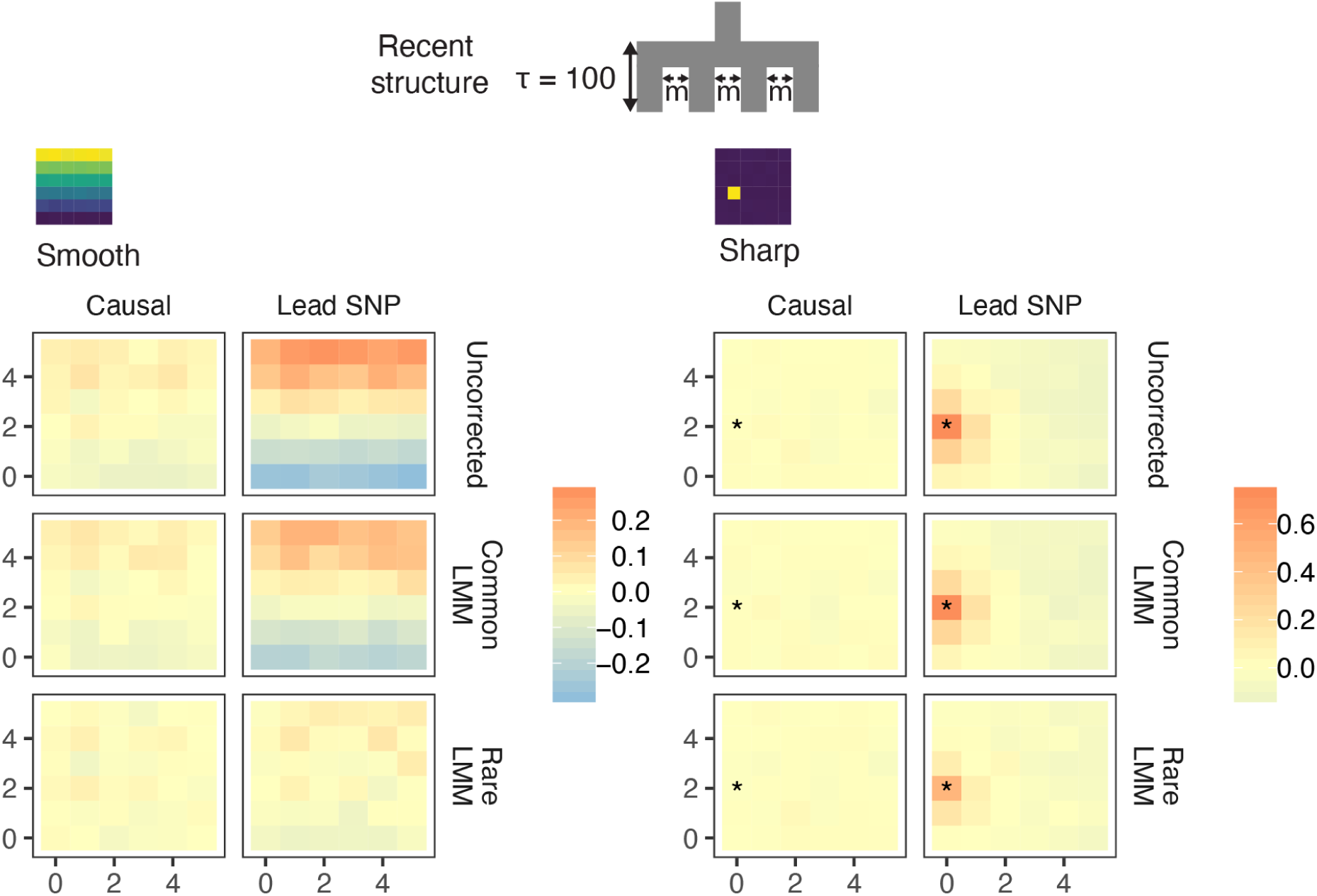
Spatial distribution of residual polygenic scores based on effect sizes from linear mixed models carried out in GCTA-LOCO. Residual polygenic score in each deme was averaged across all individuals from that deme and over 10 simulations of the phenotype. Causal and Lead SNP refers to polygenic scores constructed with known causal variants and topmost significant SNPs, respectively.

**Figure S4:**
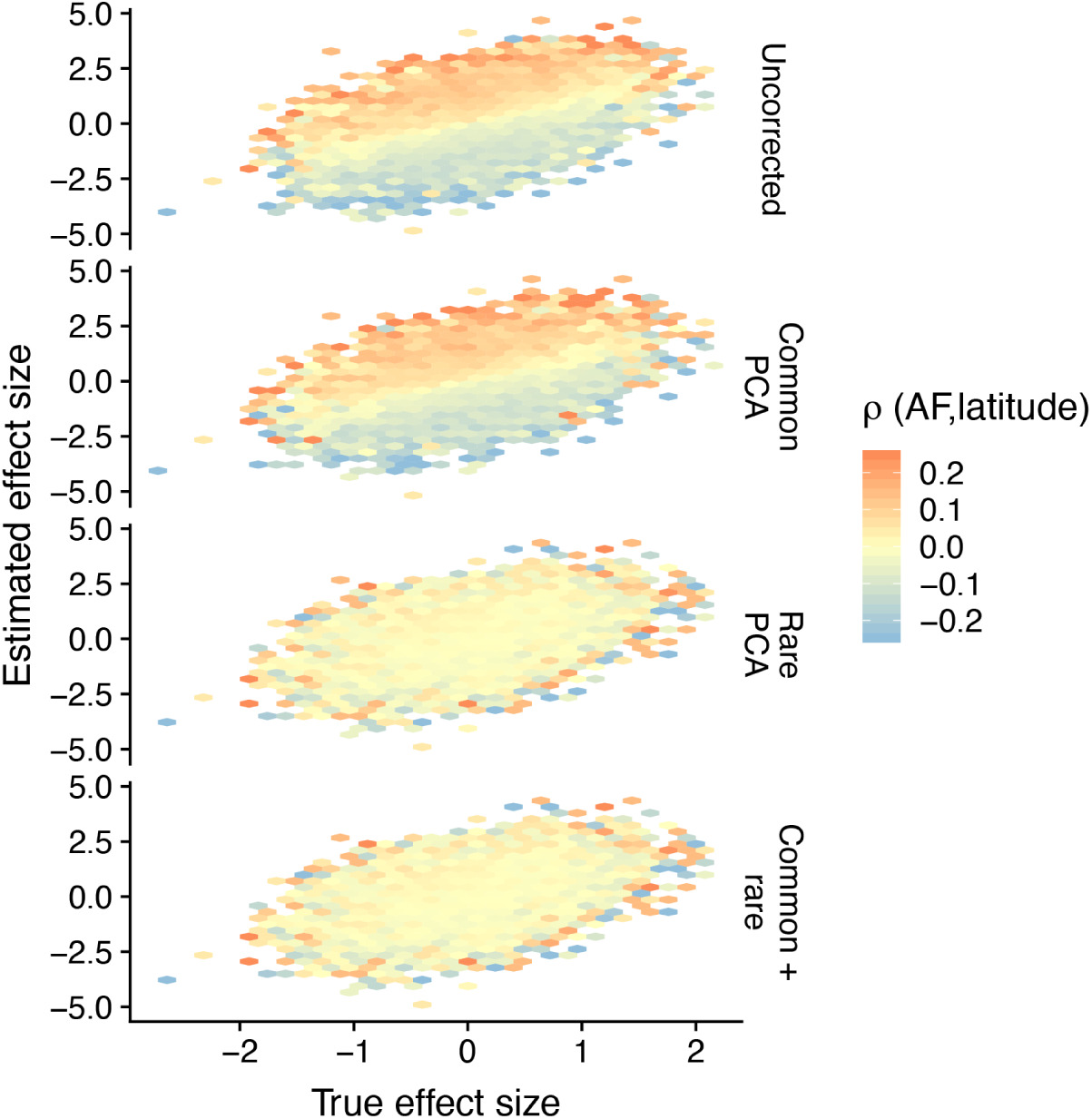
Hexagonal bin plots illustrating the residual confounding in variant effect sizes due to stratification. Each panel shows the comparison of the true (simulated) and estimated effect sizes of causal variants (M = 2,000) for different modes of correction for population structure and when the environment is distributed in a North-South cline. The color of each bin represents the mean correlation of variants in that bin and latitude, averaged across 20 simulations. When there is residual stratification, the effect sizes of variants that are positively correlated with latitude are biased upwards whereas the effect sizes of variants that are negatively correlated are biased downwards. Even though the effect size of any single variant may be biased due to stratification, the effect size across all variants is still unbiased. In this particular case, because the population structure has a recent origin, rare-PCA, but not common-PCA adequately removes the bias.

**Figure S5:**
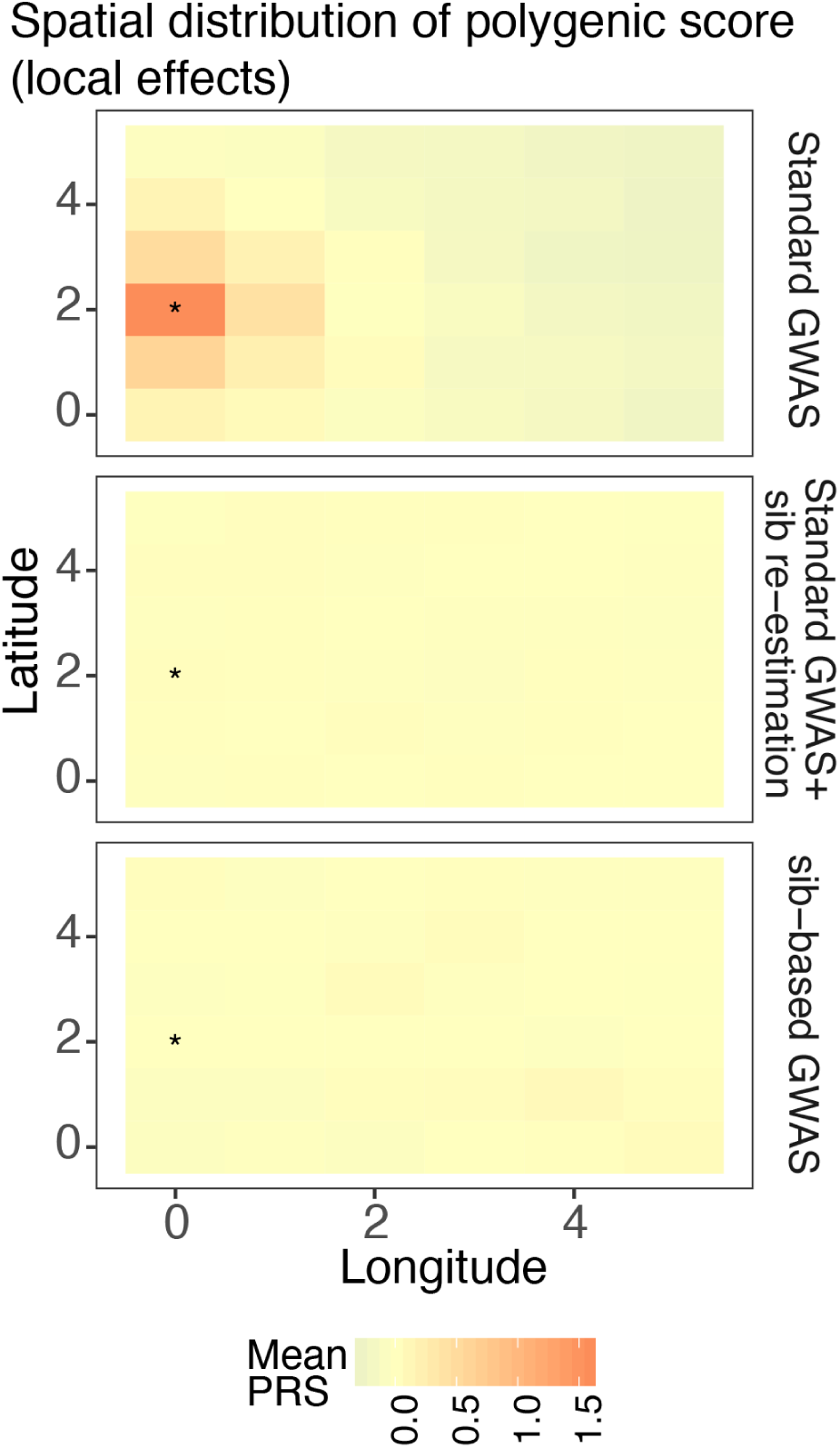
Spatial distribution of residual poygenic scores when the environment is sharply distributed (risk location indicated with *). The polygenic scores were calculated using variants and their effect sizes in a standard GWAS (top row), variants ascertained in a standard GWAS and effects re-estimated in sibling pairs (middle row), or a full sibling-based design where both variants and their effects are obtained in sibling pairs. Polygenic scores were averaged across 20 random simulations of the phenotype.

**Figure S6:**
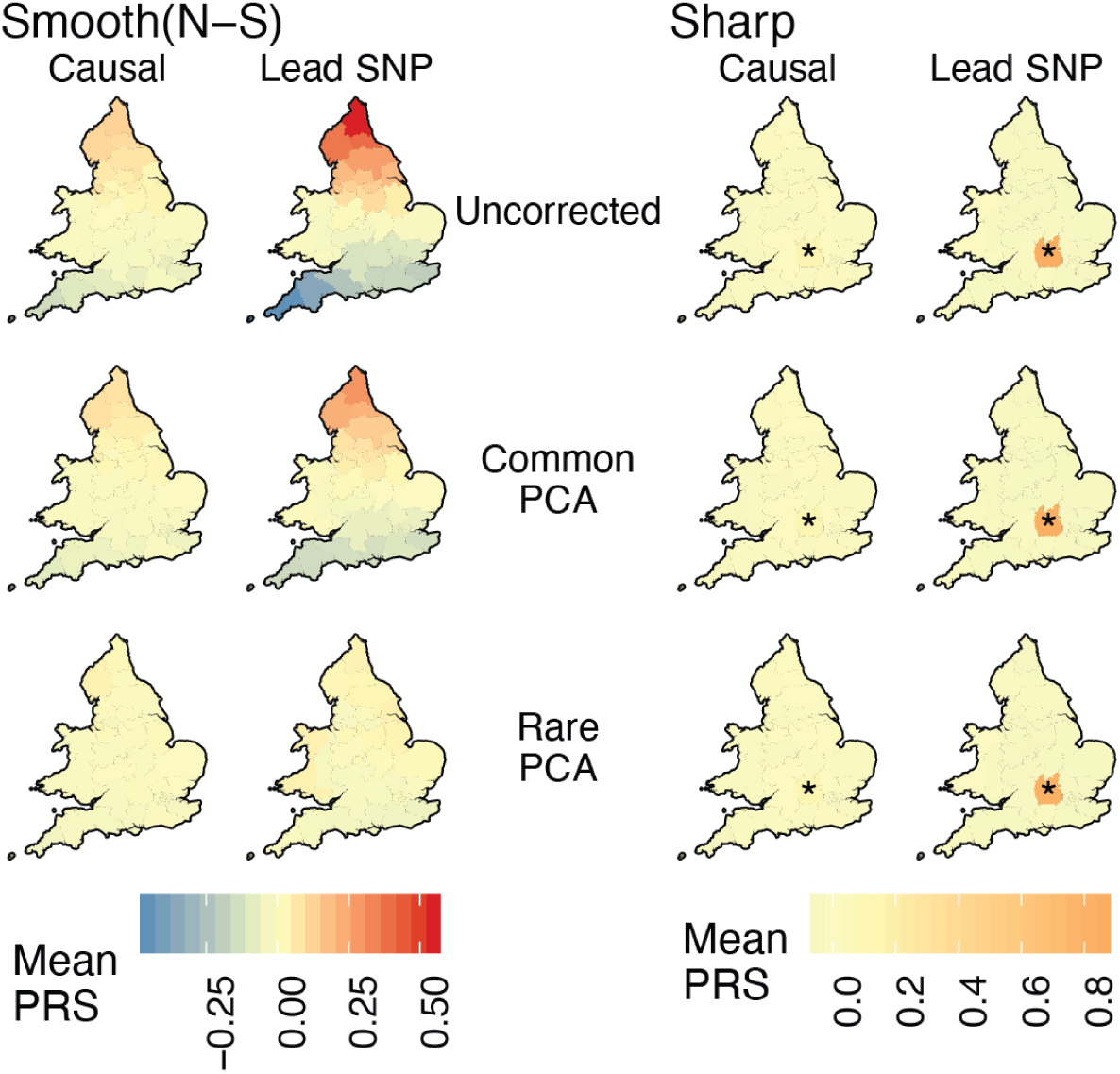
Spatial distribution of residual polygenic scores under the ‘complex’ structure model when individuals are sampled uniformly across demes. Each region is colored with the mean residual polygenic score across 250 individuals and 20 random iterations. ‘Smooth’ refers to a North-South environmental risk and ‘Sharp’ refers to a local environmental risk (risk location indicated with *). Polygenic scores were constructed using either the true causal variants (Causal) or the topmost significant SNPs (lead SNPs). Compare this to Fig.4 where individuals are sampled non-uniformly.

**Figure S7:**
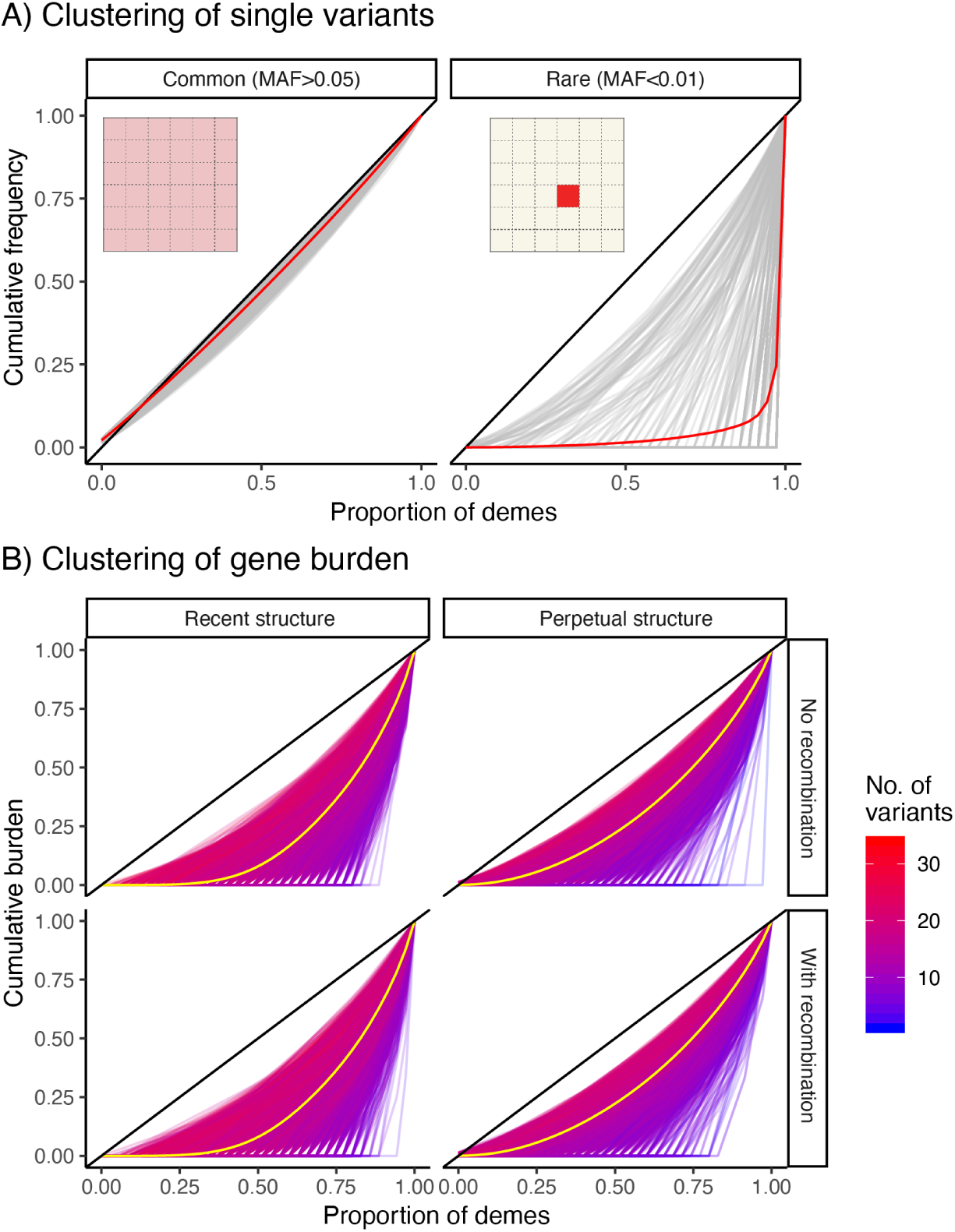
Gini curves showing geographic clustering of (A) variant frequency and (B) gene burden. (A) Grey curves are individual variants (1,000 sampled uniformly at random for plotting) and red lines indicate the mean curve across all variants. The diagonal represents the case where the variant/burden is uniformly distributed over the entire grid. Common variants tend to be more widely distributed (left panel) than rare variants (right) as illustrated by the grid inset. (B) Each curve represents a single gene (10,000 sampled) and the yellow line represents the mean across genes. The color of a curve represents the number of rare variants across which burden is aggregated. As the number of variants increases, gene burden tends to behave more like a common variant in its spatial distribution.

**Table S1:** Mean observed Fst for different migration rate under each demographic model.

| Model | Migration rate<sup>1</sup> | Mean Fst (95% C.I.) | $\lambda$ (Latitude) | $\lambda$ (Longitude) |
| --- | --- | --- | --- | --- |
| Recent | 0.001 | 3.8e-03 (3.7e-03 - 4e-03) | 3.5649 | 3.7808 |
| Recent | 0.0025 | 2.6e-03 (2.5e-03 - 2.7e-03) | 3.4733 | 3.6425 |
| Recent | 0.005 | 1.6e-03 (1.5e-03 - 1.7e-03) | 3.0914 | 3.1357 |
| Recent | 0.0075 | 1.5e-03 (1.4e-03 - 1.6e-03) | 3.4661 | 3.3344 |
| Recent | 0.01 | 1.1e-03 (1e-03 - 1.2e-03) | 3.0629 | 3.0675 |
| Recent | 0.015 | 7.9e-04 (7.2e-04 - 8.6e-04) | 2.8256 | 2.5172 |
| Recent | 0.02 | 7e-04 (6.3e-04 - 7.7e-04) | 2.4668 | 2.6838 |
| Recent | 0.025 | 5.1e-04 (4.4e-04 - 5.9e-04) | 2.2173 | 2.6485 |
| Recent | 0.03 | 4e-04 (3.3e-04 - 4.6e-04) | 2.4842 | 2.2036 |
| Recent | 0.05* | 2.3e-04 (1.7e-04 - 2.9e-04) | 1.6754 | 1.8486 |
| Perpetual | 0.06 | 2.5e-04 (1.9e-04 - 3.1e-04) | 1.8101 | 1.7606 |
| Perpetual | 0.07* | 2.0e-04 (1.4e-04 - 2.6e-04) | 1.6640 | 1.6381 |
| Perpetual | 0.08 | 1.7e-04 (1.1e-04 - 2.3e-04) | 1.5905 | 1.6658 |
| Complex | 0.05 | 3.2e-04 (2.5e-04 - 3.8e-04) | 2.6425 | 1.7480 |
| Complex | 0.06 | 2.8e-04 (2.1e-04 - 3.4e-04) | 2.1651 | 1.8637 |
| Complex | 0.07 | 2.5e-04 (1.8e-04 - 3.1e-04) | 1.9318 | 1.7012 |
| Complex | 0.08* | 1.5e-04 (9.7e-05 - 2.1e-04) | 1.6520 | 1.5214 |
| Complex | 0.09 | 1.7e-04 (1.1e-04 - 2.2e-04) | 1.6841 | 1.3892 |
| Complex | 0.1 | 1.7e-04 (1.2e-04 - 2.3e-04) | 1.5943 | 1.4719 |
| Complex | 0.12 | 1.3e-04 (7.3e-05 - 1.8e-04) | 1.4442 | 1.4395 |
| Complex | 0.15 | 7.9e-05 (2.7e-05 - 1.3e-04) | 1.2536 | 1.3123 |

<sup>1</sup> Proportion of migrants in and out of a deme per generation. Selected migration rate indicated with (\*) for each model.

